# Molecular phylogenetics illuminates the evolutionary history and hidden diversity of Australian cave crickets (Orthoptera: Rhaphidophoridae)

**DOI:** 10.1101/2024.08.11.607522

**Authors:** P. G. Beasley-Hall, S. A. Trewick, S. M. Eberhard, A. Zwick, E. H. Reed, S. J. B. Cooper, A. D. Austin

**Affiliations:** School of Biological Sciences, The University of Adelaide, Adelaide, SA, Australia; Wildlife & Ecology, Massey University Manawatū, Palmerston North, New Zealand; Subterranean Ecology Pty Ltd., Coningham, TAS, Australia; Australian National Insect Collection, CSIRO National Research Collections Australia, Canberra, ACT, Australia; Evolutionary Biology Unit, South Australian Museum, Adelaide, SA, Australia

## Abstract

Cave crickets (Orthoptera: Rhaphidophoridae) are a globally distributed group of insects found in dark, humid microhabitats including natural caves, alpine scree, and forest litter. Ten extant subfamilies are currently recognised and Macropathinae, which comprises the entirety of the fauna in South America, South Africa, Australia, and New Zealand, is thought to be the most ancient of these. The New Zealand rhaphidophorid fauna comprises high phylogenetic diversity throughout its mesic zone, with most species occurring above-ground. In contrast, the Australian fauna is poorly known, with an apparently greater relative proportion of species utilising caves as refugia. A robust phylogenetic framework is needed to underpin future taxonomic work on the group and potentially contrasting patterns of taxonomic diversity within Macropathinae. Here, we performed fossil-calibrated phylogenetic analysis using whole mitochondrial genomes and nuclear markers to reconstruct the evolutionary history of Macropathinae, with a focus on the Australian fauna. By dramatically increasing taxon sampling relative to past studies, we recovered the Australian fauna as rampantly polyphyletic, with the remaining Macropathinae nested amongst six distinct Australian lineages. Deep divergences between major clades imply additional Australian lineages remain undetected, either due to extinction or sampling bias, and have likely confounded past biogeographic signal. We inferred the radiation of Macropathinae began during the Lower Cretaceous prior to the fragmentation of Gondwana with a potential Pangaean origin for Rhaphidophoridae. Finally, we found evidence for several undescribed taxa of Australian Rhaphidophoridae, all of which qualify as short-range endemics, and discuss the conservation implications of these restricted distributions.

## Introduction

Insects are the most biodiverse group of animals on Earth and are represented by over one million described species (Stork, 2018), yet their true diversity is poorly known. Enumerating this extraordinary diversity is necessary for conservation, but this endeavour is complicated by the fact that certain species are less likely to be discovered than others. Large-bodied, geographically widespread species are more likely to be given formal scientific names, and therefore conservation attention, particularly those with visual appeal (Barua et al., 2012; Stork et al., 2008). In contrast, subterranean insects typically possess “unaesthetic” troglomorphic traits that include the elongation of sensory appendages, eye loss, and pigment reduction. These features can result in morphologically cryptic species which may only be detected through molecular phylogenetic analysis, further exacerbating a lack of taxonomic knowledge (King et al., 2022). The resulting lack of taxonomic knowledge may result in respective habitats receiving inadequate conservation effort relative to actual need. Indeed, subterranean environments—which include terrestrial and groundwater ecosystems in karstic caves, lava tubes, calcretes, and interstitial habitats—are often overlooked in conservation management efforts, despite being the largest non-marine ecosystem in the world (Mammola et al., 2022; Vaccarelli et al., 2023). Increasing taxonomic knowledge has been identified as an urgent first step towards conserving subterranean ecosystems worldwide (Wynne et al., 2021). To this end, here we use molecular phylogenetics and historical biogeography to explore the biodiversity and evolutionary history of an understudied group of Australian cave insects.

Rhaphidophoridae Walker, commonly referred to as camel or cave crickets, is a globally-distributed, largely nocturnal family of Orthoptera found on every continent except Antarctica (Cigliano et al., 2022). Rhaphidophorids are wingless and may be found in animal burrows, alpine scree, forest litter, caves, and artificial structures such as abandoned mines. Subterranean species are reliant on caves to complete their life cycles but typically forage above-ground on a nightly basis (Richards, 1966). Rhaphidophoridae presently contains 10 subfamilies, with Macropathinae Karny thought to have diversified prior to all others (Allegrucci and Sbordoni, 2019a; Dowle et al., 2024; Gorochov, 2001; Hubbell and Norton, 1978). This Gondwanan subfamily comprises the known cave cricket fauna in Australia, New Zealand, South Africa, and South America (Fig. 1). Within Macropathinae, New Zealand fauna (also called cave wētā or tokoriro) is the best studied and the most biodiverse, with most species known from forests and the alpine zone and only a handful found in caves (Fitness et al., 2018; Hegg et al., 2022, 2019; Richards, 1972; Trewick, 2024). In contrast, in Australia almost all species are predominantly recorded from caves, with above-ground aridity—and consequently less expansive wet forest habitats, especially on the mainland—implicated in determining their dependence on subterranean habitats (Richards, 1987). In the cooler, wetter island state of Tasmania, which supports extensive tracts of wet forests, aridity may be less of a factor versus the distribution of cave bearing rock types. No taxonomic research has been conducted on the Australian fauna for almost fifty years; the resulting knowledge gap is concerning because five Australian species are formally listed as threatened and human disturbance, including clearing of native forest, has been linked to population declines (Orthopteroid Specialist Group, 1996; Simms et al., 1996; Threatened Species Section, 2022a, 2022b, 2022c, 2022d).

**Fig. 1.**
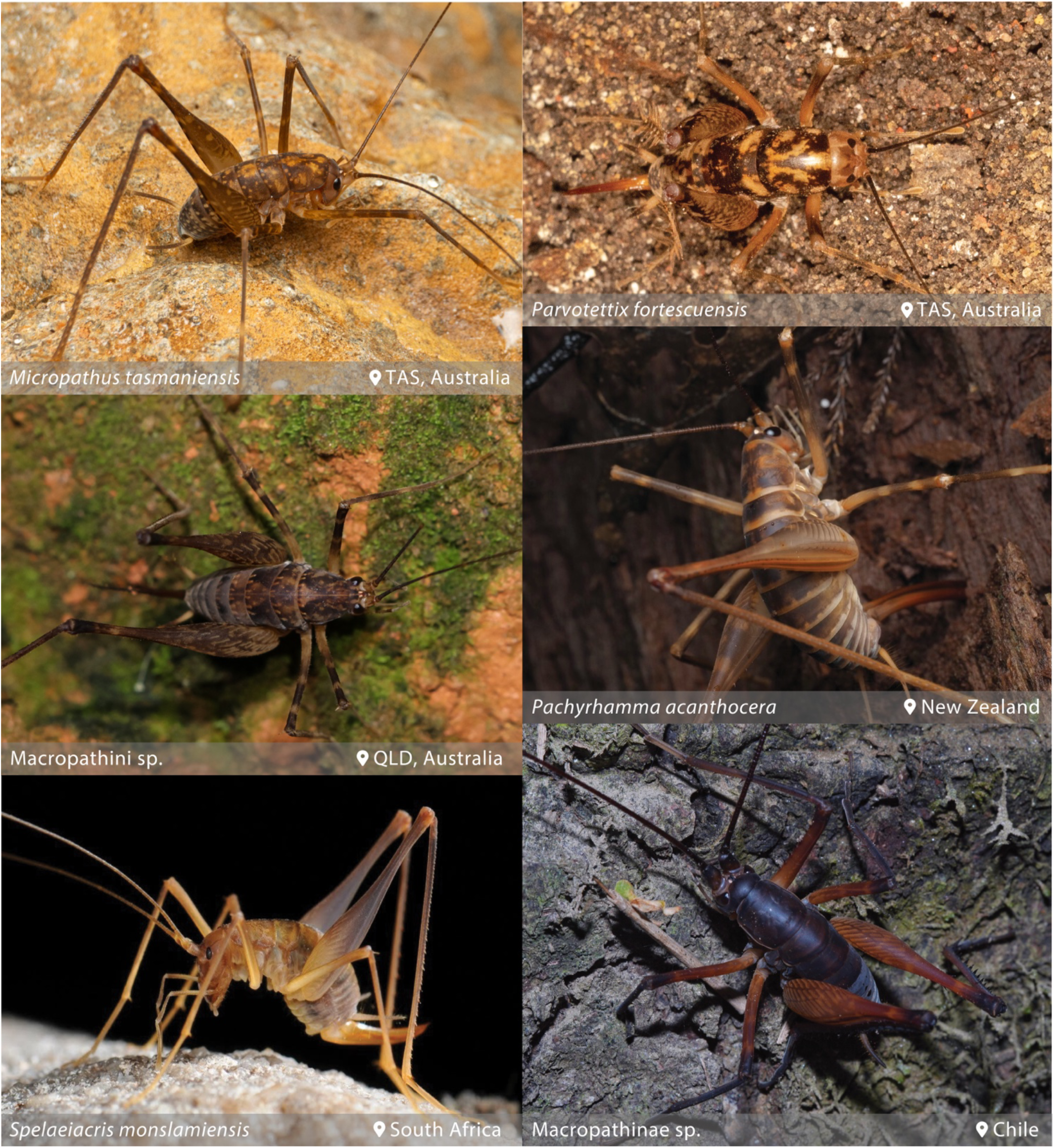
Morphological diversity of Macropathinae from throughout the Southern Hemisphere. TAS = Tasmania; QLD = Queensland. Images © Jessa Thurman, Keith Martin-Smith, Greg Tasney, commoncopper via iNaturalist, Peter Swart, and Benjamin Silva Ahumada. Photographs reproduced under Creative Commons licences.

Phylogenetic analyses have shown that the neighbouring Australian and New Zealand macropathine faunas are not reciprocally monophyletic. Examples include the Tasmanian genus *Parvotettix* Richards being the sister group to the remainder of the subfamily, including all other Australian genera (Beasley-Hall et al., 2018) and the New Zealand *Macropathus* Walker being sister to representatives from South America (Allegrucci et al., 2010). Further, recent time-calibrated phylogenetic work has highlighted several New Zealand lineages with divergence times pre-dating their island habitats, suggesting not only that a large proportion of the fauna remains to be discovered, but also that these flightless, nocturnal insects may possess a greater propensity for over-water dispersal than previously assumed (Dowle et al., 2024). Like Tasmania, high rhaphidophorid diversity in New Zealand—which has a cool, wet climate that supports wet forests—indicates that aridity, or native forest logging and clearing, reduces available habitat for these insects. The macropathine fauna of Australia therefore provides an interesting contrast in which a different pattern of taxonomic diversity might exist in cave and epigean habitats. We expect this diversity could reflect a relatively deep ancestry, echoing past mesic conditions across the continent (Byrne et al., 2011) and the utilisation of cave systems as refugia in the present day, especially in the warmer and drier regions of mainland Australia.

To paint a clearer picture of rhaphidophorid diversity in Australia, we applied molecular methods to reconstruct a comprehensive phylogeny for the group. The subfamily Macropathinae exhibits a high degree of intraspecific morphological variation (Richards, 1971a), necessitating the use of molecular data to strengthen future taxonomic inference. A robust phylogenetic framework for Australian macropathines will not only enable the discovery and description of species previously unknown to science, but also assist with documentation of genera and species that have awaited formal scientific names for decades (Richards, 1987). Here, we specifically sought to: 1) resolve relationships among the eight described Australian genera; 2) identify Australian lineages representing undescribed taxa for future taxonomic study; 3) determine relationships of the Australian fauna within the broader subfamily Macropathinae; and finally 4) enhance understanding of the phylogenetic position and biogeography of Macropathinae at the familial level.

## Materials and methods

### Taxon sampling and DNA sequencing

Australian cave cricket specimens were obtained from museum collections and in the field (see Fig. S1 for sampling map). Museum specimens were sourced from the South Australian Museum (SAMA), the Tasmanian Museum and Art Gallery (TMAG), the Queensland Museum (QM), the Australian Museum (AM), the Melbourne Museum (MM), and the Australian National Insect Collection (ANIC). We sampled all 22 described species of Australian Rhaphidophoridae with the exception of *Speleotettix chopardi* (Karny), which represents a *nomen dubium* (Beasley-Hall *et al.* submitted), as well as several putatively undescribed species and genera. We also integrated published GenBank data from six other rhaphidophorid subfamilies to infer the placement of Macropathinae at the family level, namely Aemodogryllinae Jacobson and Rhaphidophorinae Walker (East and Southeast Asia), Dolichopodainae Brunner von Wattenwyl and Troglophilinae Karny (Mediterranean), and Ceuthophilinae Tepper and Tropidischiinae Scudder (North America). Additional GenBank data for Prophalangopsidae Kirby (grigs), Anostostomatidae Saussure (king crickets), Tettigoniidae Krauss (katydids), and Caelifera Ander (grasshoppers and allies) were used as outgroups to facilitate fossil calibration and root the tree. A complete list of taxa included in the present study, as well as relevant citations and accession numbers, is available in Supplementary Table S1.

For all specimens except those housed at ANIC (see below), DNA was extracted from mesothoracic leg tissue using a Gentra Puregene Tissue Kit (QIAGEN). The mitochondrial genes *12S* rRNA, *16S* rRNA, and cytochrome oxidase I (*COI*) and nuclear *28S* rRNA (D1 and D3-5 regions) were amplified using the following primers: 12Sai + 12Sbi (*12S*) (Simon et al., 1994), 16sar + 16sbr (*16S*) (Cunningham et al., 1992), LCO1490 + HCO2198 (*COI*) (Folmer et al., 1994), 686 + 427 (*28S* D1 region) (Friedrich and Tautz, 1997), and 485 + 689 (*28S* D3-5 regions) (Friedrich and Tautz, 1997). PCRs were performed using a standard hot-start protocol consisting of a preheat step at 95ºC for 10 minutes, denaturation at 95ºC for 45 seconds, annealing at 48ºC (*COI, 28S* D1), 51ºC (*12S* and *16S* rRNAs), or 55ºC (*28S* D3-5) for 45 sec, extension at 72ºC for 1 min, 72ºC for 10 min, and 25ºC for 2 min 20 sec. Denaturation, annealing, and extension steps were repeated for 39 cycles. Bi-directional Sanger sequencing was outsourced to the Australian Genome Research Facility (AGRF, Melbourne, Australia) under standard PCR cycling conditions.

Sanger sequences were processed and aligned using the package *sangeranalyseR* in R v.4.3.0 (Chao et al., 2021). Sequences were trimmed by quality scores using a Phred score cutoff of 0.001 (Q30) and modified Mott trimming. A signal ratio cutoff of 0.33 and minimum fraction call of 0.75 were selected for base calling. Next-generation sequencing data were processed using the Trim Galore wrapper (github.com/FelixKrueger/TrimGalore) for Cutadapt and FastQC (Andrews, 2010; Martin, 2011) for quality trimming, filtering, and adapter removal. Quality-controlled reads were then *de novo* assembled using SPAdes with k-mer lengths of 33, 55, 77, 91, and 121, and the identity of resulting contigs was verified using a MegaBLAST search (Camacho et al., 2009) against the nucleotide collection of the GenBank repository (Sayers et al., 2023). For specimens with smaller contigs requiring scaffolding against a reference, reads were mapped against the closest available relative in GenBank with the *Map to Reference* tool in Geneious Prime v.2023.1.2 (geneious.com) using medium sensitivity presets and without trimming before mapping.

We conducted next-generation sequencing (NGS) on two subsets of our specimens. The first consisted of type specimens housed at ANIC, sequenced as part of an existing project run by the Commonwealth Scientific and Industrial Research Organisation (CSIRO). ANIC specimens were extracted in-house using a modified Qiagen DNeasy Blood and Tissue kit protocol in 384-well format and sequenced using a miniaturised genome skimming approach centred around the Qiagen QiaSeq UltraLow Input library kit. Pools of 384 libraries were sequenced on an Illumina NovaSeq S1 flow cell. Our second subset was selected after initial analysis to provide whole mitogenome data for at least one specimen per major clade, as well as additional nuclear markers (*18S* rRNA and *histone-3* [H3]) to strengthen node support at the backbone of our tree. This subset also included museum specimens that did not successfully PCR amplify for Sanger sequencing. Library preparation was performed by AGRF using the Illumina DNA Prep (M) (fresh specimens) or IDT xGen Library Prep workflow (museum specimens). Paired-end sequencing was conducted by AGRF on an Illumina NovaSeq 6000 or NovaSeq X Plus platform, respectively, to generate 150-base-pair reads. Coding regions, tRNAs, and rRNAs in resulting mitogenome sequences were annotated using Prokka v.1.14.6 (Seemann, 2014). Annotated mitogenomes were processed for GenBank submission with GB2sequin (Lehwark and Greiner, 2019).

### Phylogenetic analysis

Alignments of mitochondrial protein-coding genes, mitochondrial rRNAs, *H3, 18S*, and two regions of *28S* (aligned separately) were produced using MUSCLE v.3.8.425 (Edgar, 2004) with default settings. Alignments were then concatenated and divided into nine partitions: mitochondrial protein-coding genes separated by codon position (1, 2, and 3), mitochondrial rRNAs (*12S*+*16S* together), *H3* separated by codon position (1, 2, and 3), *18S* rRNA, and finally *28S* rRNA (D1+D3-5 regions together) for a total of 18,146 base pairs. The resulting molecular matrix used for downstream is available in Supplementary File S1.

We used maximum likelihood (ML) and Bayesian inference (BI) for phylogenetic analysis and divergence dating. ML analysis was performed in IQ-TREE v.1.6.12 (Nguyen et al., 2015) using ModelFinder for substitution model selection (*-m MFP+MERGE*) and 1000 ultrafast bootstrap replicates (Hoang et al., 2018; Kalyaanamoorthy et al., 2017). We performed Bayesian inference using BEAST2 v.2.6.4 (Bouckaert et al., 2019). We selected the package bModelTest for substitution model selection (Bouckaert and Drummond, 2017), the birth-death tree prior—suitable for datasets containing inter- and intraspecific data (Ritchie et al., 2017)— and the optimised relaxed clock (Douglas et al., 2021). To assist the analysis through the burn-in stage, we set starting clock rates of 1.0e-6 and 1.0e-9 for mitochondrial and nuclear genes, respectively (Bouckaert, 2015). We ran three separate chains of 100 million generations with a tree logging frequency of every 10,000 generations. Convergence of the stationary distribution and effective sample size values (>200) of parameters were assessed using Tracer v.1.7.2 (Rambaut et al., 2018). Log files and trees were combined in LogCombiner v.2.7.5 with a 10% burn-in. A maximum clade credibility tree was inferred using TreeAnnotator v.2.7.1 (Bouckaert et al., 2019) with the common ancestors approach to node height annotation and a posterior probability limit of 0.5.

### Divergence dating

To date the timings of cladogenesis within Rhaphidophoridae, we constrained our BI analysis with our ML topology as a starting tree such that only branch lengths were inferred throughout the run. We chose to do this because our initial BI analysis (described above) recovered the South American fauna as polyphyletic with poor node support, whereas it was inferred as a well-supported monophyletic group in our ML analysis in agreement with several previous studies (Allegrucci and Sbordoni, 2019b; Dowle et al., 2024; Kim et al., 2024). As such, we had reason to believe our ML tree reflected a more plausible set of relationships within Rhaphidophoridae. To constrain our BI analysis, we first scaled the ML starting tree using treePL to ensure compatibility with our chosen fossil calibration (Smith and O’Meara, 2012). Our dated Bayesian analysis was run in BEAST2 as for the uncalibrated analysis but only required three chains to reach convergence.

While several well-dated rhaphidophorid fossils exist, namely †Protroglophilinae Gorochov from the late Eocene, their phylogenetic placement remains unresolved (Allegrucci and Sbordoni, 2019a; Gorochov, 2001) and there are therefore no appropriate ingroup fossils to calibrate the molecular clock for the family. As such, we calibrated the node leading to Prophalangopsidae, Tettigoniidae, and Anostostomatidae using †*Pseudaboilus wealdensis* Gorochov, Jarzembowski & Coram, which has been rigorously assessed for its appropriateness in calibrating the orthopteran molecular clock and used in a recent phylogenomic analysis of the order (Song et al., 2020). This fossil has been dated to a minimum age of 127.5 Mya (Barremian, Lower Cretaceous) and is conservatively considered stem-(Tettigonioidea Krauss+Hagloidea Handlirsch) (Gorochov et al., 2006; Song et al., 2020). We applied the calibration using an exponential distribution with 127.5 Mya as a hard minimum bound and 271.8 Mya, the minimum age estimate of the oldest definitive Ensifera Chopard (Bethoux et al., 2002; Wolfe et al., 2016), as a soft maximum.

### Historical biogeography

We used BioGeoBEARS (Matzke, 2014, 2013) in R v.4.4.0 (R Core Team, 2024), a package using probabilistic models for ancestral range estimation, to visualise historical biogeographic hypotheses for Macropathinae and relatives. Species distributions were compiled from observations made by Richards (Richards, 1987) and the Orthoptera Species File as at June 2024 (Cigliano et al., 2022). Species were coded as belonging to Australia (including Tasmania), New Zealand (including subantarctic islands), South America, South Africa, Southeast Asia, the Mediterranean, and/or North America. Historical biogeographic analyses were run using three models: dispersal-extinction-cladogenesis (DEC) (Ronquist, 1997), a ML implementation of dispersal-vicariance analysis (DIVALIKE) (Ree and Smith, 2008), and Bayesian Analysis of Biogeography (BAYAREALIKE) (Landis et al., 2013). Each of these models assumes a different set of possible sympatric and/or vicariant processes associated with cladogenetic events (Matzke, 2013). Founder-event speciation can also be incorporated into BioGeoBEARS models, but we chose not to use this parameter (*j*) because our primary goal was to estimate ancestral ranges rather than the mode of speciation or diversification. We ran these models with default parameters using our time-calibrated phylogeny and their relative probabilities were assessed using the Akaike (AIC) and corrected Aikaike (AICc) information criteria.

## Results

### Phylogenetic analysis

We recovered mitochondrial (*12S* rRNA, *16S* rRNA, and *COI*; whole mitogenomes where possible) and nuclear (*18S* rRNA, two subunits of *28S* rRNA, and *H3*) molecular markers from 131 representatives of Australian Rhaphidophoridae. For samples for which we recovered whole mitogenomes, up to 18,094 bp were sequenced; for those represented by multi-locus Sanger data only, we recovered up to 5,966 bp. Using ML and BI, we recovered a monophyletic Macropathinae consisting of six major clades (Figs. 2, 3). Sister to all other macropathines in the analysis was *Parvotettix* and an undescribed genus, the most recent common ancestor (MRCA) of which diverged from the rest of the subfamily *ca.* 119.1 Mya (95% highest posterior density [HPD] 88.91–159.87 Mya). The MRCA of the South African genus *Spelaeiacris* Péringuey diverged *ca*. 99.78 Mya (95% HPD 71.78–136.75 Mya), and a second Australian lineage emerged *ca.* 85.16 Mya (95% HPD 62.53–115.59 Mya) containing the southeast Australian *Speleotettix* Chopard and *Cavernotettix* Richards. A widespread group then arose *ca*. 67.97 Mya (95% HPD 56.9–105.02 Mya) containing the New Zealand and South African faunas as well as two unrelated Australian lineages, the Tasmanian *Tasmanoplectron* Richards and Queensland *Australotettix* and relatives. The Tasmanian *Micropathus* Richards diverged from the MRCA of the southern Australian *Pallidotettix* Richards and *Novotettix* Richards *ca.* 63.74 Mya (95% HPD 46–87.98 Mya), which themselves diverged from one another *ca*. 31.55 Mya (95% HPD 21.39– 44.14 Mya). Aemodogryllinae and Rhaphidophorinae, representing the Indomalayan realm in our analysis, were recovered as the sister clade to Macropathinae. Sister to this broader group was a Laurasian clade containing Dolichopodainae, Troglophilinae, Ceuthophilinae, and Tropidischiinae.

**Fig. 2.**
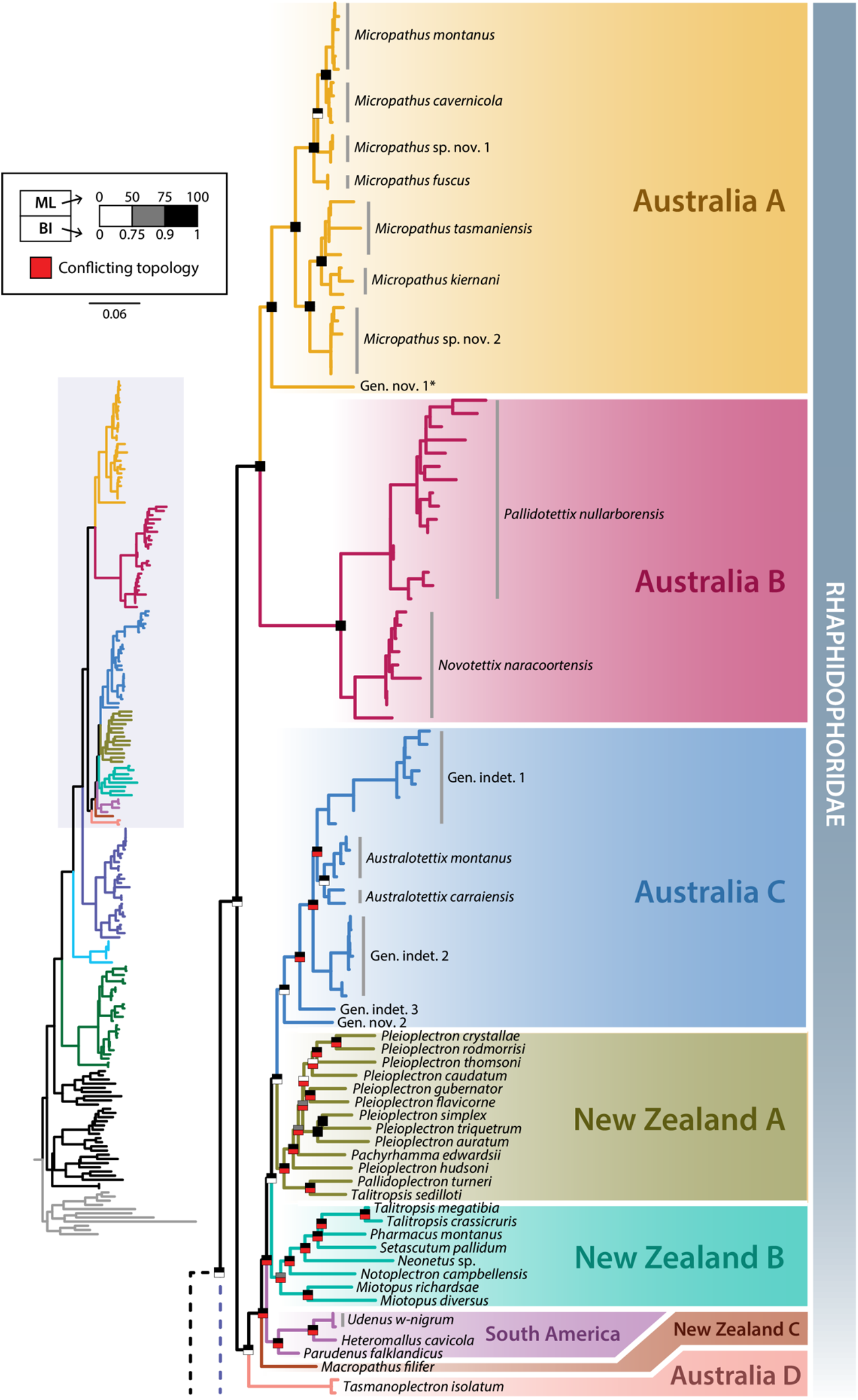
Maximum likelihood phylogeny of Macropathinae (part) inferred from mitochondrial and nuclear data. *Described as *Eburnocauda* while the present study was in review (Iannello and Beasley-Hall, 2024). Conflicting topologies refer to disagreement between maximum likelihood and Bayesian analyses. Phylogeny continued in Fig. 3. A map of sampling locations is available in Fig. S1.

**Fig. 3.**
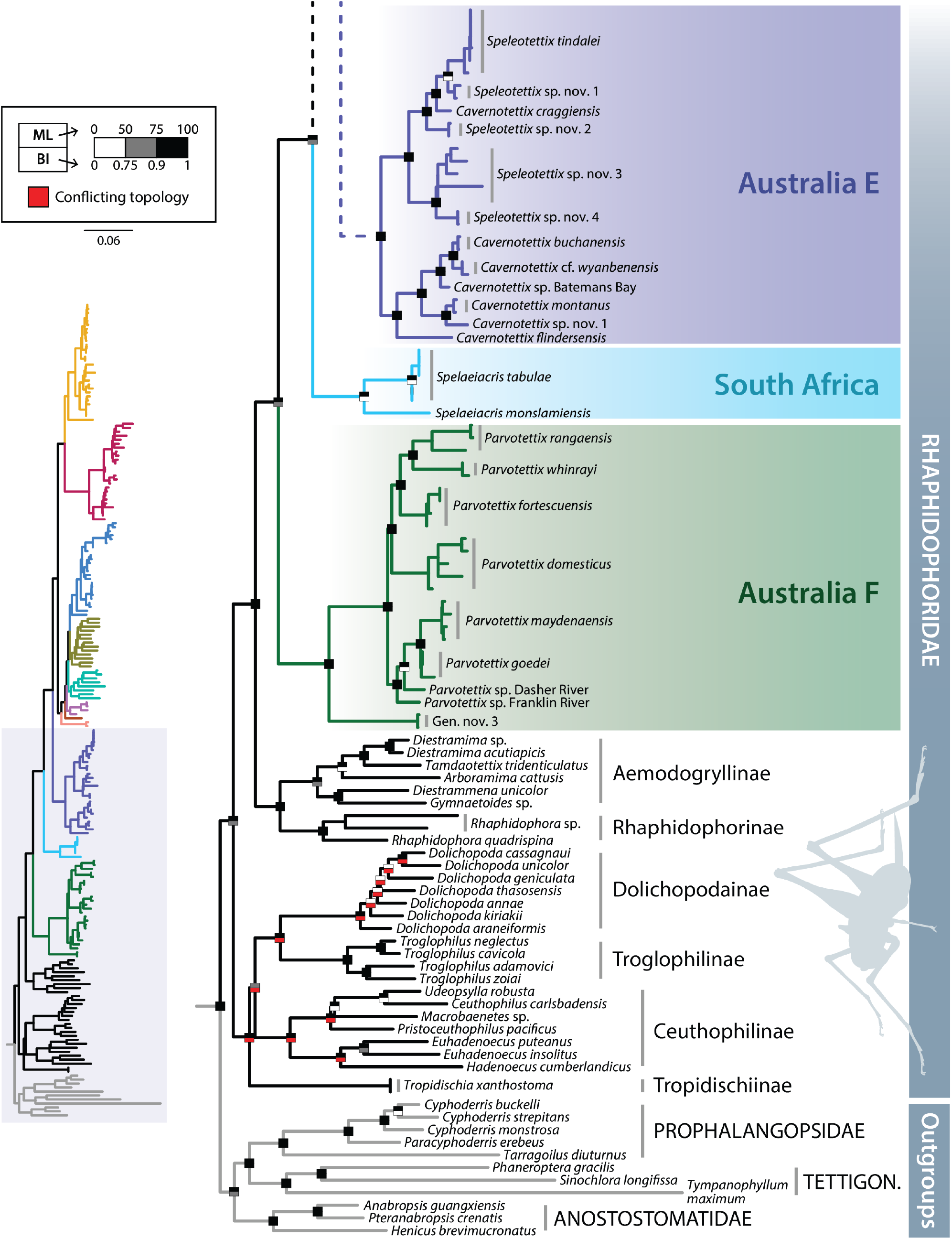
Phylogeny of Macropathinae (part; coloured branches) and related subfamilies of Rhaphidophoridae (black branches) inferred from mitochondrial and nuclear data, continued from Fig. 2. Conflicting topologies refer to disagreement between maximum likelihood and Bayesian analyses. TETTIGON. = Tettigoniidae. A map of sampling locations is available in Fig. S1. The caeliferan outgroup used in the analysis is not shown for clarity.

Several Australian lineages were not recovered as members of described species and/or genera and represent putatively new taxa (Figs. 2–3). These include two species of *Micropathus* (clade Australia A); a new genus sister to *Micropathus* (Australia A); three lineages corresponding to new species of *Australotettix* Richards, if not new genera in their own right (labelled indeterminate genera [gen. indet.] for this reason; Australia C); a new genus related to *Australotettix* (Australia C); four species of *Speleotettix* (Australia E); one species of *Cavernotettix* (Australia E); and a new genus sister to *Parvotettix* (Australia F). Additional lineages were suggestive of new species within existing genera but were only represented by one sample, leaving their identifications less certain (e.g., *Cavernotettix* sp. from Batemans Bay and *Parvotettix* spp. from Franklin and Dasher Rivers, Figs. 2–3).

Our ML and BI analyses recovered similar relationships among major clades of Macropathinae and with high node support in most cases. However, they differed with respect to two clades (Figs. 2, 3). In our ML analysis, the New Zealand fauna (except for *Macropathus filifer* Walker) was inferred as sister to northeastern Australian species (Australia C) and *Macropathus* was sister to a monophyletic South American clade (New Zealand C, Fig. 2). In contrast, an initial BI phylogeny recovered a non-monophyletic South American fauna interspersed by New Zealand species but with low node support, prompting us to constrain the topology of our downstream time-calibrated BI tree. Interestingly, the same ML/BI conflict was reported by (Kim et al., 2024) for the New Zealand and South American Rhaphidophoridae. The position of Tropidischiinae also differed between our ML and BI analyses (Figs. 2, 3), but the Laurasian taxa were nonetheless recovered as monophyletic and relationships among non-macropathine subfamilies were not the main focus of the present study. Raw output files from IQTREE and BEAST2 are available in the electronic supplementary material (Supplementary Files S2, S3).

### Historical biogeography

The DEC model was the best fit to our dataset based on AIC and AICc scores (AIC = 78.82414; AICc = 79.02086) compared to DIVALIKE (86.22179; 86.41851) and BAYAREALIKE (435.22473; 435.42146) (Fig. S2). In historical biogeographic analysis, a *range* refers to the geographic range of a taxon and can consist of one or more *areas*. BioGeoBEARS infers the relative probability of different ancestral ranges; here and below we refer to most-probable ancestral ranges. In our analysis, Australia, New Zealand, South America, and South Africa (i.e., Gondwana) was estimated as the ancestral range of the subfamily (Fig. S2). At the root of our tree, Rhaphidophoridae was estimated to have a broad ancestral range spanning all areas except the Mediterranean.

## Discussion

Here we present a comprehensive molecular phylogenetic analysis of Rhaphidophoridae with a focus on Macropathinae, a subfamily representing the majority of Southern Hemisphere biodiversity. Our results expand on previous findings demonstrating the Australian and New Zealand faunas are both non-monophyletic, with the former thought to consist of at least two distantly related clades based on limited sampling (Allegrucci and Sbordoni, 2019a; Beasley-Hall et al., 2018). In the current study, we recovered Macropathinae as monophyletic and containing a diverse, polyphyletic Australian fauna consisting of at least six major lineages, with the remainder of the Southern Hemisphere fauna nested between or within those groups. We recovered the New Zealand fauna as similarly polyphyletic and sharing a most recent common ancestor with South American species (Fig. 2), in agreement with a recent phylogenetic study focusing on the Rhaphidophoridae of New Zealand (Dowle et al., 2024).

We were primarily interested in examining the relationships and biogeography of Macropathinae in the present study, but we also included additional representatives of the family to assess its placement within the broader Rhaphidophoridae. Our findings are inconsistent with morphology-driven hypotheses that proposed Macropathinae as the most “primitive” subfamily within Rhaphidophoridae and sister to all others (Ander, 1939; Gorochov, 2001; Hubbell and Norton, 1978; Karny, 1930). Instead, our results agree with other phylogenetic studies showing Macropathinae is most closely related to Aemodogryllinae and Rhaphidophorinae from Southeast and Eastern Asia (Figs. 4, S2) (Allegrucci and Sbordoni, 2019a; Dowle et al., 2024; Kim et al., 2024). However, these data do collectively suggest the subfamily diversified prior to other major groups of Rhaphidophoridae, supporting its “ancient”—albeit not “primitive” or arguably basal—status.

**Fig. 4.**
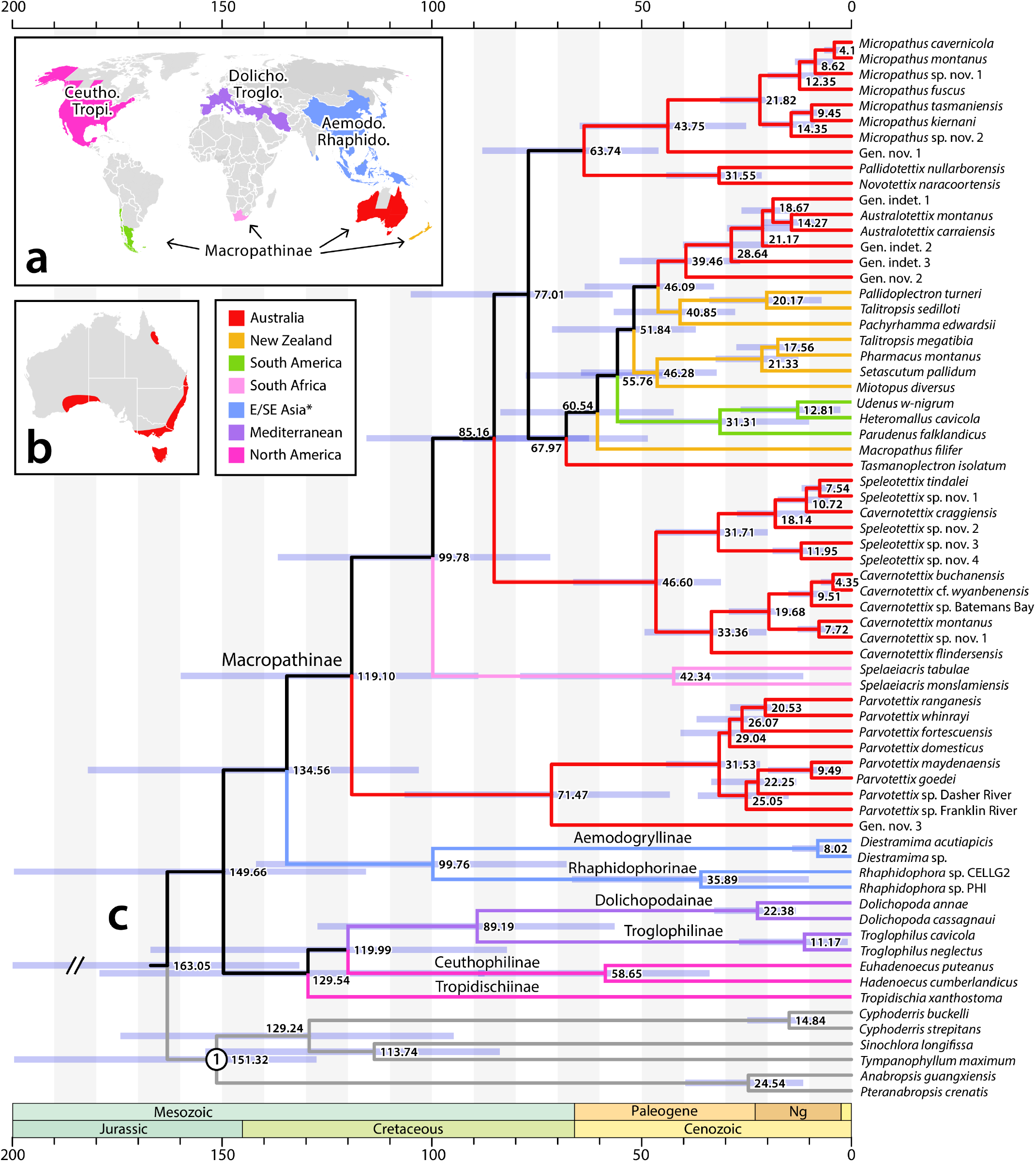
a) Distribution of rhaphidophorid subfamilies examined here. b) Known distribution of Macropathinae in Australia. c) Time-calibrated Bayesian phylogeny of Rhaphidophoridae using a single fossil calibration (node 1). Estimated timings of cladogenesis are shown at nodes; node bars indicate 95% HPD interval ranges. *Aemodogryllinae is also found in East Asia, but only Southeast Asian species were included in our analysis.

### Biogeography of Macropathinae

Previous age estimates for Rhaphidophoridae have largely been derived from order-level studies that have not extensively sampled the family, with a tendency to focus on a handful of Northern Hemisphere taxa (Song et al., 2020; Vandergast et al., 2017). Two recent studies have expanded sampling of Rhaphidophoridae to more closely examine its age and diversification. Kim *et al.* (2024) focused largely on the Southeast Asian and Mediterranean Rhaphidophoridae and constrained their tree using secondary age estimates from a fossil-calibrated phylogeny by Song *et al.* (2020). Dowle et al. (2024) focused on the New Zealand Macropathinae and used a fossil and geological calibration to estimate the age of the subfamily. While the age estimates of Dowle *et al.* (2024) were slightly older than Kim *et al*. (2024) and the present study, all three estimated an age of around 140 Mya for the most recent common ancestor of Macropathinae. As for previous studies, this order and timing of diversification is consistent with the family originating prior to the separation of Pangaea, with major cladogenetic events reflecting the separation of the supercontinent into Laurasia and Gondwana (Allegrucci and Sbordoni, 2019a; Kim et al., 2024).

Our results support the hypothesis that the macropathine fauna radiated prior to the fragmentation of Gondwana, with the earliest representatives appearing on the Australian landmass. Firstly, the South African, South American, and New Zealand faunas are all nested between, or within, Australian clades in the phylogeny (Figs. 2, 3). Secondly, we infer the MRCA of Macropathinae diverged from the remainder of Rhaphidophoridae in the Lower Cretaceous 134.56 Mya (95% HPD 103.11–182 Mya), well before the separation of Australia from Antarctica and New Zealand *ca*. 75 Mya (Fig. 4) (McIntyre et al., 2017). Not all cladogenetic events in our tree reflect Gondwanan breakup: for example, based on median age estimates the MRCA of the South African *Spelaeiacris* diverged from other Macropathinae *ca*. 99.78 Mya, placing it after the separation of Africa from West Gondwana *ca*. 110 Mya (McIntyre et al., 2017). Long distance dispersal via rafting has been proposed to resolve this apparent contradiction between inferred divergence times and geological events, e.g., in the case of New Zealand wētā (Dowle et al., 2024). At the family level, we estimated a late Jurassic age of *ca.* 149.66 Mya (95% HPD 115.68–199.63 Mya) for Rhaphidophoridae.

While continental vicariance and dispersal have likely both influenced the present-day distribution of Rhaphidophoridae, additional factors must be considered when the biogeographic history of the group. The diverse Australian fauna is separated by deep divergences in our phylogeny, even when only minimum ages of nodes are considered (Fig. 5), suggesting that more closely related lineages have not been sampled. For example, we inferred several island species (*Cavernotettix craggiensis* Richards, *flindersensis* (Chopard), and sp. nov. 1; *Parvotettix rangaensis* Richards and *whinrayi* Richards) diverged from their closest relatives millions of years before the isolation of such land masses over the past *ca.* 14,000 years due to post-glacial flooding (Lambeck and Chappell, 2001). This observation of “old” taxa on “young” islands has also been documented for the New Zealand Rhaphidophoridae (Dowle et al., 2024) and discussed throughout the broader phylogenetic literature (McCulloch and Waters, 2019). To explain this pattern, Dowle *et al.* (2024) suggested the interplay between speciation and extinction processes in Rhaphidophoridae have resulted in a mismatch between geological history and biology. In other words, biogeographic signal is confounded due to a failure to sample (e.g., non-island) sister species, either because those lineages are extinct or as-yet unknown to science. A combination of both may have led to similar patterns in the Australian fauna in our phylogeny. In addition to extinction processes resulting from extensive aridification and Pleistocene climatic changes to the Australian continent (Byrne et al., 2008; Weij et al., 2024), the cryptic nature of individuals in both forest and subterranean habitats has almost certainly contributed to incomplete sampling of rhaphidophorid fauna globally. This is especially applicable in an Australian context given the apparent high abundance of the group in caves, disjunct distributions of species in relictual pockets of wet forest, and the inaccessibility of such locations for sampling. Failure to sample for this reason would imply a greater diversity in Australia than currently known, a scenario supported by the discovery of several undescribed taxa in our phylogeny even with modest sampling.

#### Conservation and taxonomic implications

Rhaphidophorids are an important faunal component of caves in the Australian mesic zone where they play a prominent ecological role as predators, omnivorous scavengers, and detritivores. The abundance of rhaphidophorid biomass also supports predator communities (Richards, 1971b) and they are the key food source sustaining populations of the iconic Tasmanian cave spider *Hickmania troglodytes* (Higgins & Petterd) (Driessen, 2009). Further, all described Australian cave cricket species apparently have exceptionally limited distributions (<10,000 km^2^), qualifying them as short-range endemics (SREs) at a higher risk of extinction compared to more cosmopolitan taxa (Harvey, 2002; Richards, 1971a). Despite this, only 68 of a described 879 species globally have had some form of conservation assessment per the IUCN Red List; over half of those assessed are considered at risk of extinction (Cigliano et al., 2022; IUCN, 2023). As a result, establishing which species are undescribed *vs*. a disjunct population of a more widespread taxon—as we have done here—is a key step in the management of these SREs and the development of clear conservation priorities.

Our findings have several conservation implications for the Australian Rhaphidophoridae. Five species are currently represented in federal or state threatened species lists: *Cavernotettix craggiensis, Micropathus kiernani* Richards, *Parvotettix rangaensis, Parvotettix whinrayi*, and *Tasmanoplectron isolatum* Richards, all found in Tasmania (Orthopteroid Specialist Group, 1996; Threatened Species Section, 2022a, 2022b, 2022c, 2022d). As all described Australian species fulfil the criteria for SREs, many more are likely in need of conservation listing. Key threatening processes include native forest logging and land clearing, which risk fragmenting wet forest habitat providing above-ground food sources and corridors to connect cave populations; predation from invasive species, such as rats; and more direct human disturbance via recreational and tourist activities in caves (IUCN, 2023; Richards, 1987; Simms et al., 1996). Here, we have revealed a far higher diversity in Australia than the eight genera and 22 species currently known, highlighting at least three new genera and 15 new species (Figs. 2, 3). “Gen. nov. 1” in Fig. 2 was described as the monotypic genus *Eburnocauda* Beasley-Hall & Iannello while the present study was in review, bringing the total number of rhaphidophorid genera in Australia to nine (Iannello and Beasley-Hall, 2024). Ultimately, our findings have increased the known biodiversity of Rhaphidophoridae in Australia by at least 50% but a far greater species richness almost certainly remains undiscovered.

## Conclusions

We have presented a robust phylogeny of cave crickets (Rhaphidophoridae) with a focus on the Southern Hemisphere subfamily Macropathinae. By substantially expanding upon past sampling of the Australian fauna, we have confidently resolved relationships among all described genera. We have also identified at least 15 new species and three new genera based on molecular data, which will form the basis of future taxonomic work. We found the Australian fauna is rampantly polyphyletic and consists of five major unrelated lineages, three of which have independently emerged in Tasmania. We have also demonstrated that the New Zealand and South African Rhaphidophoridae share ancestors with the Australian fauna, and that South American cave crickets are in turn nested within a polyphyletic New Zealand fauna. We estimate ancestral Macropathinae appeared *ca*. 134 Mya, diverging from a common ancestor shared with the Asian Rhaphidophorinae and Aemodogryllinae, and began to diversify across Gondwana in the Lower Cretaceous *ca*. 119 Mya. We infer the Rhaphidophoridae itself originated *ca*. 150 Mya across Pangaea.

Improving understandings of evolutionary relationships and taxonomy within Rhaphidophoridae is important for several reasons. First, cave crickets represent a valuable, yet often cryptic, ecological component of subterranean realms. Identifying species boundaries is necessary for conservation management of these short-range endemics, with particular relevance to the understudied Australian biota. Second, robust phylogenetic frameworks can assist in identifying characters that are informative for future taxonomic research and identification by non-scientist stakeholders (e.g., park managers, naturalists, and speleologists). Finally, organismal groups like Rhaphidophoridae can shed further light on the link between changing climatic conditions, species distributions, and lineage diversity. For example, by comparing the Australian and New Zealand fauna in the present study, we have highlighted that cave crickets are not simply a “relictual” fauna only found underground—as is often the narrative associated with Northern Hemisphere species—but can be highly diverse in cool, wet forests, with many lineages still awaiting taxonomic description even in the relatively restricted mesic zone of Australia.

## Acknowledgements

The authors have no conflict of interest to disclose. This work was funded by a National Taxonomy Research Grant Program fellowship awarded to PGBH by the Australian Government Department of Climate Change, Energy, the Environment and Water (no. 4-H3JJWE), with co-funding from the Australian Speleological Federation, South Australian Museum, Environment Institute, and The University of Adelaide. We would like to heartily thank the staff, curators, and collection managers of institutions around Australia for sending us material: Simon Grove and Kirrily More (Tasmanian Museum and Art Gallery); You Ning Su and Federica Turco (Australian National Insect Collection); Matthew Shaw and Ben Parslow (South Australian Museum); and Karin Koch (Queensland Museum). Finally, we acknowledge The University of Adelaide’s Phoenix High Performance Computing service for contributing resources for the phylogenetic analyses presented here.

Specimens were collected under the following permits held by PGBH unless otherwise stated: AA-0001076 via Parks Victoria and FS/14-3694/1_2024 via the Victorian Department of Energy, Environment and Climate Action (Victoria); U26509-1 and Y27154-2 via the South Australian Department for Environment and Water, the latter held by ER (South Australia); SL102683 via the New South Wales Department of Planning and Environment, held by the Australian Museum (New South Wales); and TFA 22452 via the Tasmanian Department of Natural Resources and Environment, held by SME (Tasmania).

## Data availability

*Please note: the project ID and URLs below will be updated to citable, persistent DOIs upon acceptance of this manuscript.*

All raw data associated with this study are available via FigShare via project ID XXXXX. Briefly, Supplementary Table S1 contains GenBank accession numbers for molecular sequences and associated specimen metadata (https://figshare.com/s/a3bc3cd0f98b3a5c4037); Supplementary File S1 contains the molecular matrix used as input for phylogenetic analyses in NEXUS format (https://figshare.com/s/fd5dc6ab3e31f033ad56); and Supplementary Files S2 and S3 contain raw phylogenetic trees from the maximum likelihood (https://figshare.com/s/7a83846a1cfc2ad586ba) and time-calibrated Bayesian phylogenetic analyses (https://figshare.com/s/41a9d8ccc82136eed1f6), respectively.

