## Supplementary Material for "Molecular phylogenetics illuminates the evolutionary history and hidden diversity of Australian cave crickets (Orthoptera: Rhaphidophoridae)"

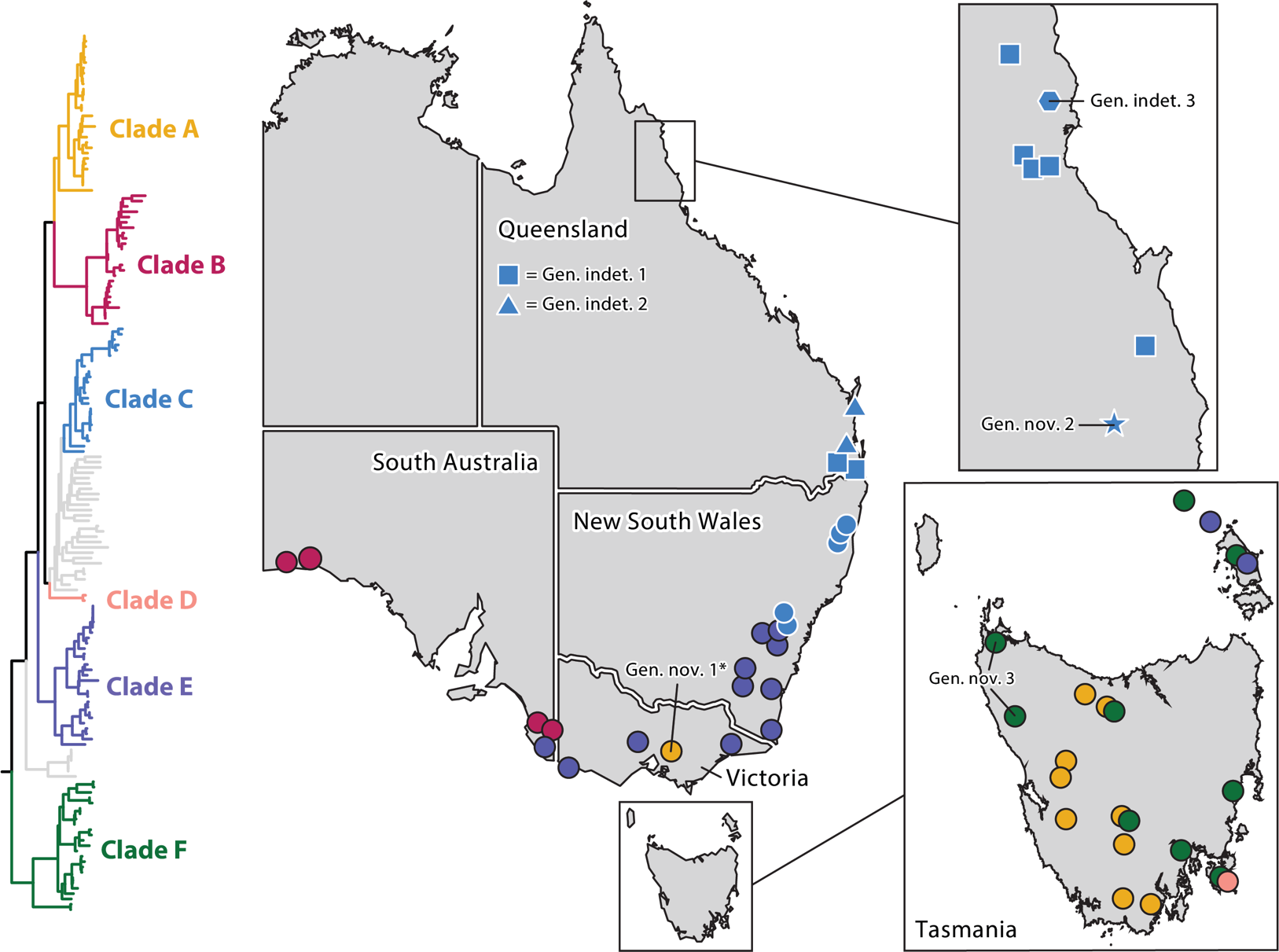

**Fig. S1.** Locations sampled in the present study. Populations are colour-coded by clade and undescribed taxa highlighted. *Described as *Eburnocauda* in Iannello and Beasley-Hall (2024) while the present study was in review.

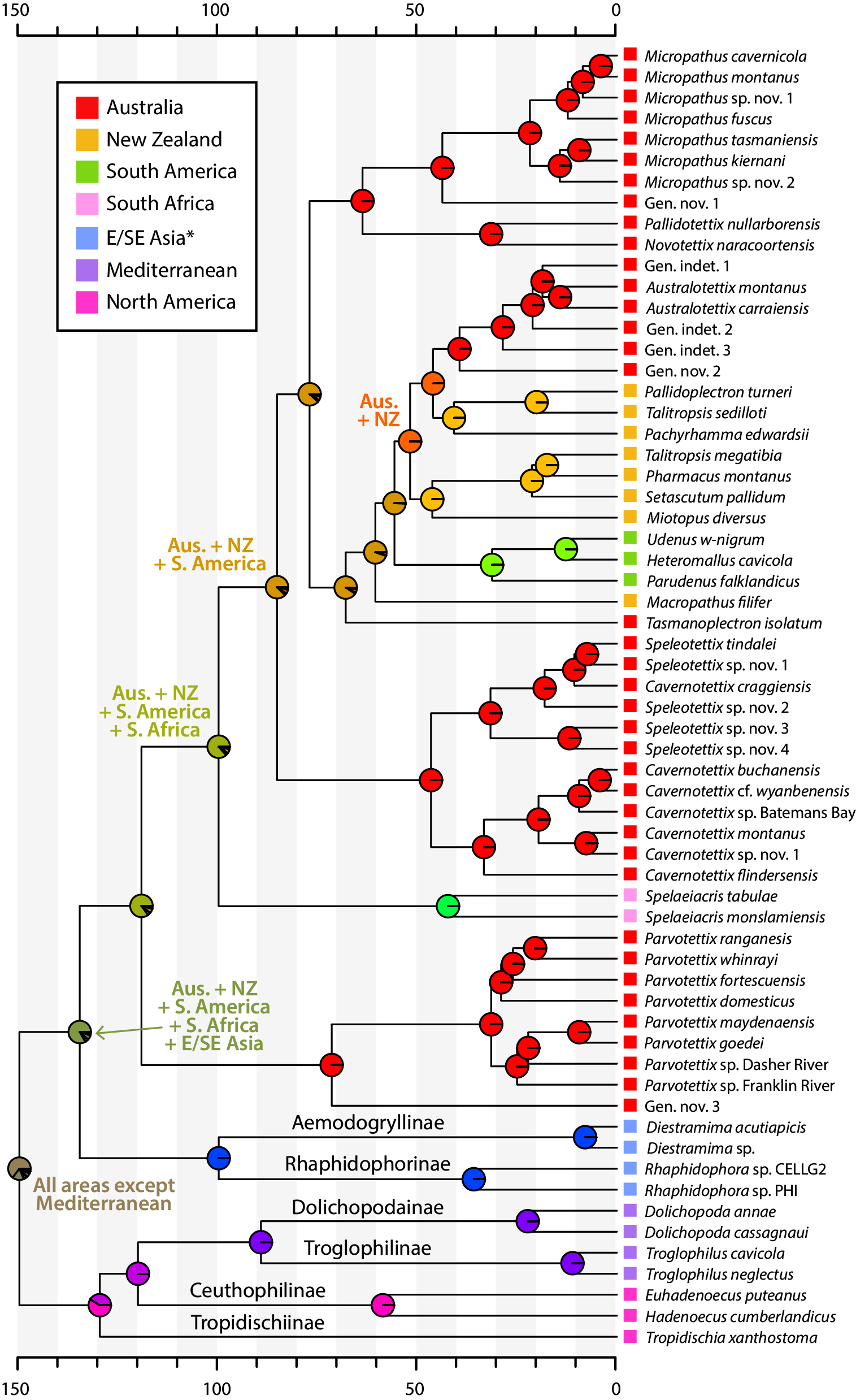

**Fig. S2.** Ancestral areas of Rhaphidophoridae estimated using the DEC model in BioGeoBEARS. Pie slices indicate the relative probability of different ranges instantaneously before cladogenesis. Contemporary distributions are shown at tips. *Aemodogryllinae is also found in East Asia, but only Southeast Asian species were included in our analysis. The caeliferan outgroup used in the analysis is not shown for clarity.

***Please note: FigShare URLs provided below are private links and will be updated to persistent, citable DOIs upon acceptance of this publication.***

**Table S1.** Rhaphidophorid specimens included in the present study and their associated collection and institution metadata. GenBank accessions of DNA sequences are coloured by whether they were generated here (black) or retrieved from existing studies (grey). Available via FigShare: <https://figshare.com/s/a3bc3cd0f98b3a5c4037>

**File S1.** Molecular matrix, in NEXUS format, used as input for maximum likelihood and Bayesian phylogenetic analysis. Partitions are specified in the character block at the end of the file. Available via FigShare: <https://figshare.com/s/fd5dc6ab3e31f033ad56>

**File S2.** Raw .tree file from the maximum likelihood phylogenetic analysis conducted with IQ-TREE. Available via FigShare: <https://figshare.com/s/7a83846a1cfc2ad586ba>

**File S3.** Raw .tree file from the time-calibrated Bayesian phylogenetic analysis conducted with BEAST2. Available via FigShare: <https://figshare.com/s/41a9d8ccc82136eed1f6>
